# An optimized protocol for the creation of constructs expressing sodium channel subunits to be used in functional studies of genetic epilepsies

**DOI:** 10.1101/2023.11.08.565006

**Authors:** Saúl Lindo-Samanamud, Amanda M. do Canto, Fabio R. Torres, Marina C. Gonsales, Alexandre B. Godoi, Douglas C. da Rosa, André S. Vieira, Iscia Lopes-Cendes

**Affiliations:** Department of Translational Medicine, School of Medical Sciences, University of Campinas (UNICAMP), Campinas, Brazil; The Brazilian Institute of Neuroscience and Neurotechnology (BRAINN), Campinas,Brazil; Department of Structural and Functional Biology, Institute of Biology, University of Campinas (UNICAMP), Campinas, Brazil

**Keywords:** NEB Stable competent cells, plasmids, transformation, mutagenesis, *SCN1A* gene

## Abstract

**Background:** We aim to develop a method for cloning highly unstable plasmids encoding sodium channel subunits expressed in the brain.

**Method:** NEB Stable competent cells (NSCC) were transformed with the native plasmid. After purification, plasmids with the expected profile of enzymatic digestion were sequenced. Finally, mutant linear plasmids (MLP) created by site-directed mutagenesis were used to transform NSCC.

**Results:** We found that choosing a suitable host for transformation and subcloning, the controlled temperature during cell growth, and a small volume of Luria Bertani (LB) medium allowed the generation of plasmids without undesirable mutations or structural alterations.

**Comparison with existing method:** At present, there is little information on a detailed method for cloning plasmids carrying the SCN1A gene. Using the method proposed here, we observed increased cell growth and precision in the sequence of the obtained plasmids.

**Conclusion:** We have developed an optimized strategy for replicating unstable plasmids using NSCC with an appropriate volume of culture medium and an adequate temperature in the cloning and subcloning phases. We have also demonstrated how to create the desired plasmid constructs by using site-directed mutagenesis. We expect that our protocol will assist investigators interested in functional studies of sodium channels expressed in the brain.

## 1. INTRODUCTION

Bacterial cells can be transformed, incorporating new genetic traits by acquiring a physiological state called “competence” (Veening and Blokesch, 2017). This state allows the expression of genes related to the absorption and processing of DNA. When exogenous DNA is incorporated into the cytoplasm, it can integrate into the bacteria’s genome or become a plasmid (Etchuuya et al., 2011; Million-Weaver and Camps, 2014). Plasmids are circular extrachromosomal DNA fragments with the ability to self-replicate and generate new phenotypes, allowing the host cell to adapt to environmental changes (Boucher et al., 2013; Ridenhour and Top, 2016).

Recombinant DNA technology allows plasmid engineering in *Escherichia coli*. However, several technical issues must be overcome to maintain and replicate the plasmid correctly (Al-Allaf et al., 2013). The main challenges are related to the size and constitution of the plasmid DNA, the technique used to introduce the exogenous DNA, the bacterial strain used in the transformation, the competence of the bacterial cell, and the cell growth protocol used (Corchero and Villaverde, 1998; Yoshida and Sato, 2009). If these parameters are not optimized, plasmid DNA stability and quality, as well as the expression of the cloned product, could be affected negatively. The possible causes for failure in these experiments are that the cell cannot maintain an adequate plasmid synthesis metabolism, or it may be affected by the stress of the culture maintenance. In addition, intrinsic characteristics of the host bacterium could lead to plasmid instability, causing unwanted losses or alterations in the plasmid sequence (Silva et al., 2012).

Voltage-dependent sodium channels (VGSC) are transmembrane proteins that allow sodium ions to flow across the cell membrane. These channels are hetero-oligomeric polypeptides formed by a functional alpha unit associated with one or two modulatory beta subunits. One of these alpha units is the Nav1.1 protein encoded by the *SCN1A* gene, which is located on chromosome 2q24.3. *SCN1A* has 26 exons (Wallace et al., 2001), and mutations in *SCN1A* are a major cause of monogenic forms of epilepsy (Meng et al., 2015).

One alternative to study the functional effect of mutations in VGSC is by expressing the mutation in a heterologous system allowing for cell electrophysiological studies (Lossin et al., 2002; Sendfeld et al., 2019). However, these heterologous expression systems containing plasmids encoding subunits of sodium channels require complex and laborious cloning processes, a factor that frequently results in experimental failures. Furthermore, recent research has revealed the presence of sequences in the cDNA of the *SCN1A* gene that resemble prokaryotic promoters. These sequences lead to the transcription of mRNAs that are detrimental for translation in bacteria. This could potentially contribute to the high instability observed in these plasmids. This emphasizes the importance of implementing optimized protocols in related studies(DeKeyser et al., 2021). In this context, we report here a novel and optimized protocol using the NEB Stable bacteria (New England Biolabs) as an efficient host for plasmids containing the wild-type and mutant forms of the *SCN1A* gene. In addition, we describe in detail how to generate and confirm mutations using site-directed mutagenesis and next-generation sequencing technology.

### 2. MATERIAL AND METHODS

### 2.1. Plasmids and bacterial strains

We used the pCMV-*SCN1A* plasmid, which is derived from the vector pCMV-script. The vector is linked to a 6230 bp fragment that includes the coding sequence and the 3′ untranslated region (UTR) of the Nav1.1 channel (*SCN1A* gene); the size of the recombinant plasmid is 10451 bp (Lossin et al., 2002). The plasmid carrying the *SCN1A* gene cDNA was given by Dr. Alfred George Jr. (Department of Pharmacology, Northwestern University Feinberg School of Medicine, Chicago, Illinois, USA). Plasmid cloning was carried out using two competent *E. coli* strains: the NEB Stable competent cells (NSCC, New England Biolabs) and Stbl2 cells (Thermo Fisher Scientific). To better characterize the pCMV-*SCN1A* plasmid, we performed an *in-silico* analysis to identify the repetitive units and the regions more susceptible to recombination, using the REPFIND programs (https://zlab.bu.edu/mfrith/cgi-bin/repfind.cgi).

### 2.2. Transformation with wild type plasmids

We kept the native plasmid pCMV-*SCN1A* on ice for 10 minutes in a sterile 1.5-ml microtube. We then immediately added 25 μl of the NSCC, avoiding bubbles, and left it on ice for 30 minutes. Subsequently, we moved the microtube containing the plasmid and cells to a water bath at 42°C for 30 seconds and immediately transferred the microtube to the ice for 2 minutes. After that, we added 400 μl of NEB® 10-beta/Stable Outgrowth Medium and placed the microtube in the incubator at 30°C for 90 minutes at 225 rpm. This is the bacterial replication stage; therefore, it is important to control the incubation temperature, which should be between 27 and 30°C, to optimize the growth of the colonies. Subsequently, we added 113 μl of the inoculum to four Petri dishes (90 × 15 mm), containing 25ml solid Luria Bertani (LB) medium with 12,5 μl of kanamycin (50 mg/ml). We spread this inoculum avoiding the borders of the LB plate and let the plate dry for 3 minutes, turned it upside down, and transferred it to the incubator at 27°C for 2–3 days.

### 2.3. Subcloning

After 48 hours, cell colonies were identified, isolated by using the tip of a sterile toothpick, and grown in sterile 1.5-ml microtubes containing 1200 μl of liquid LB medium with 0,6 μl of kanamycin (50 mg/ml). Microtube caps were perforated to promote gas exchange. All steps were performed in a sterile laminar flow cabinet. The microtubes were transferred to an oven at 27°C and 225 rpm for one day when cell cultures became turbid.

### 2.4. Extraction and purification

We used the commercially available Gen Elute Plasmid Miniprep kit (Sigma) according to the following protocol. Overnight bacterial cultures (600 μl) were centrifuged at 12000 *g* for 1 minute at 27°C.The remaining 600 μl of inoculum was mixed with 600 μl of 50% glycerol and stored at -20 °C. Cell pellets were processed according to the manufacturer’s protocol. These plasmids were stored at -20°C for their subsequent characterization.

### 2.5. Quantification and analysis of the enzymatic profile

The purified plasmid was quantified by fluorometry and diluted to a concentration of 30 ng/μl in a final volume of 20 μl. Plasmids were subjected to enzymatic digestion to evaluate the presence of structural rearrangements (enzymatic digestion profile). The enzyme used was FastDigest BamHI (Thermo Scientific), which digests the plasmid pCMV-*SCN1A* to generate a digestion profile of four DNA fragments (Positive Profile). We performed restriction digestion by using 17 μl of the plasmid DNA (30 ng/μl), 2 μl of FastDigest Green Buffer, and 1 μl of the restriction enzyme BamHI. We incubated this reaction in a temperature cycler at 37°C for 5 minutes. The digested plasmids were then subjected to electrophoresis in a 0.8% agarose gel at 100 V and 300 mA for 35 minutes and then visualized in an ultraviolet radiation transilluminator. We made a culture on LB-Agar, we took a miniculture with glycerol frozen at -80ºC, with a bacteriological seeding loop of 10 μL and we spread the content on the LB-Agar plate supplemented with Kanamycin. We let it grow for 24 hours at 30ºC inside an oven. After the colony has grown, we chop a colony with a sterile filter tip and seed into the Falcon with 4mL of LB-Broth supplemented with Kanamycin, incubate for 24 h at 250rpm at 30ºC. Then we seeded 400 uL of the culture to an Erlenmeyer with 200mL of LB-Broth supplemented with kanamycin and left at 250rpm at 30ºC/ for 16h.The purification and extraction of the plasmids were carried out with the GenElute ™ HP Plasmid Midiprep kit (Sigma).

### 2.6. Next generation sequencing of plasmids

According to the manufacturer’s standard protocol, libraries were prepared using NimbleGen Seqcap Ez library SR version 4.3 (Roche) and sequenced on a Miseq (Illumina) benchtop sequencer. We performed plasmid fragmentation by sonication using an M220 Focused ultrasonicator (Covaris) following these parameters: power (50 W), temperature (20°C), cycle per base (200 cycles), duty factor (20%), and time (255 seconds). The KAPA kit (Roche) was adopted for constructing the plasmid library, which uses SeqCap EZ Choice custom capture panels. The library pool was quantified by real-time PCR (7500 Real-Time PCR System, Applied Biosystems) and diluted to 4 nM with the elution buffer containing Tween 20 (EBT). Next, we denatured the library pool using NaOH and added a hybridization buffer (HT1) to dilute the library to a final concentration of 12 pM. Samples were submitted to next generation sequencing by using a nanoFlowcell V2. All subsequent steps were performed in a Miseq benchtop sequencer, including cluster generation and 2 × 150 paired-end sequencings. Finally, we performed data analysis by using our in-house open-source pipeline.

### 2.7. Selection of variants and *in silico* prediction analysis

The variants selected for site-directed mutagenesis were previously reported in a cohort of Dravet syndrome (DS) patients (Gonsales et al., 2019). Three of them were found *de novo* and are considered implicated in the etiology of the disease. However, the other variant was also found in the unaffected mother and controls, suggesting non-pathogenicity.

To estimate the deleterious effects of the selected variants, we used 13 prediction algorithms: FATHMM^*^, Condel^†^, MutationTaster^‡^, PANTHER^§^, SNPs&GO^**^, MutPred2^††^, PROVEAN^‡‡^, CADD^§§^, PolyPhen2^***^, MutationAssessor^†††^, SIFT^‡‡‡^, Align GVGD^§§§^, and PhD-SNP^****^.

### 2.8. Site-directed mutagenesis

Mutations were introduced into the plasmid using the GeneArt® Site-Directed Mutagenesis PLUS Kit (Thermo Fisher Scientific) and pCMV-*SCN1A* as the template plasmid. Mutagenesis primers were designed with the Primer X tool (http://www.bioinformatics.org/primerx). Four different mutation types were introduced in exon 26 of the *SCN1A* gene, resulting in four mutant plasmids. The synthesis of these mutations was performed in two steps: standardization of the primers to create the mutations and creation of the mutant plasmids.

#### 2.8.1. Standardization of the primers

A mixture of the forward and reverse primers (both at 10 mM) was used to generate the mutations of interest. We used a polymerase with high processivity and fidelity (AccuPrime Pfx DNA Polymerase, Thermo Fisher Scientific) for PCR reactions. Each mutation was standardized individually. The PCR master mix was prepared according to the indications of the manufacturer of the mutagenesis kit (Thermo Fisher Scientific): We mixed 95 μl of master mix containing 82.2 μl of PCR-grade water, 10 μl of 10X AccuPrime™ Pfx Reaction mix, 2 μl of the native parental plasmid (20 ng/μl), and 0.8 μl of AccuPrime ™ Pfx (2.5 U/μl). This master mix was aliquoted into five PCR tubes with 19 μl of the master mix per tube. Then, we added 1 μl of the primer mixture. The mutations were located in exon 26 of the *SCN1A* gene. They were generated by using the following PCR amplification protocol: initial denaturation of 94°C for 2 minutes; 18 cycles of denaturation at 94°C for 2 seconds, 57°C for 30 seconds, and 68°C for 5 minutes and 30 seconds; and a final extension of 68°C for 5 minutes.

PCR products (linear plasmid carrying the mutation) were subjected to 0.8% agarose gel electrophoresis at 100 V and 300 mA for 35 minutes. In addition, amplicons with the expected size (10.5 kb) were sequenced by NGS, following the same protocol described in section 2.6.

#### 2.8.2. Creation of the mutant plasmids

After PCR confirmed the generated mutations, we performed site-directed mutagenesis, which has three steps.

1. *Methylation reaction*: We methylated the parental plasmid pCMV-SCN1A using the GeneArt® Site-Directed Mutagenesis PLUS Kit (Thermo Fisher Scientific) this was performed following the manufacturer’s instructions. NSCC transformation was then performed (according to the protocol described in section 2.2) using 4 μl of the methylation reaction containing the parental plasmid. In addition, an unmethylated parental plasmid was used as a positive transformation control.
2. *Parental plasmid amplification reaction*: To create the variants of interest, the methylated parental plasmid was amplified by PCR, the standard conditions of which are described in section 2.8.1. Electrophoresis was performed on a 0.8% agarose gel to verify whether the amplicons were 10.5 kb in size. PCR products with the appropriate size were stored at -20oC
3. *Creation of SCN1A gene variants*: The 10.5 kb PCR products are clones of the parental plasmid; however, they are linear and need to be recombined with a recombinant enzyme provided by the GeneArt® Site-Directed Mutagenesis PLUS Kit (Thermo Fisher Scientific) this was performed following the manufacturer’s instructions. We performed the recombination reaction to one group of PCR products, and these recombinant products were immediately used in the transformation of both NSCC and Stbl2 cells to assess the metabolic capacity of the strains on the plasmid. Another set of PCR products did not undergo recombination; we used linear PCR products. These were used in the transformation of both bacterial strains to determine which strain has the best ability to recirculate and replicate the plasmid (metabolic capacity of the strains).
4. *Next generation sequencing of PCR products*: The PCR products were sequenced to identify whether the mutation of interest is present after mutagenesis. Libraries were prepared by using NimbleGen Seqcap Ez version 4.3 SR library (Roche) according to the protocol described above and sequenced on a Miseq benchtop sequencer (Illumina).

### 2.9. Transformation with mutant plasmids

The NSCC and Stbl2 *E. coli* strains were transformed with the recirculated mutant plasmids. Subsequently, the extraction, purification, plasmid restriction enzyme profile analysis, and next generation sequencing were performed following the abovementioned protocols. In addition, NSCC cultures carrying mutant plasmids were submitted to purification with the GenElute™ HP Plasmid Midiprep kit (Sigma). Recovered plasmids were stored at -20° C.

## 3. RESULTS AND DISCUSSION

### 3.1. Plasmids

The native plasmid pCMV-*SCN1A* was designed to express the alpha subunit of the Nav1.1 channel in eukaryotic cells. The expression is controlled by the human cytomegalovirus (CMV) promoter, which allows constitutive expression of genes cloned into the pCMV-script vector. *In silico* analysis (Figure 1) identified frequent repetitive units composed mainly of guanines (G) and cytosines (C) in the sequence of the native plasmid pCMV-*SCN1A*.Therefore, we found the presence of three clusters of CGC triplets between nucleotides 7072 and 10451 (P = 3.33067e-16). These correspond to the promoter region for the *SCN1A* gene, the resistance gene for kanamycin (*kanR*), and the plasmid origin of replication. The presence of these repetitive regions in the promoter region might lead to mutations that could disrupt the expression of the Nav1.1 protein in the cell membrane. In addition, mutations in the origin of replication or the *kanR* gene could inactivate cell replication, resulting in the absence of bacterial colonies.

**Figure 1.**
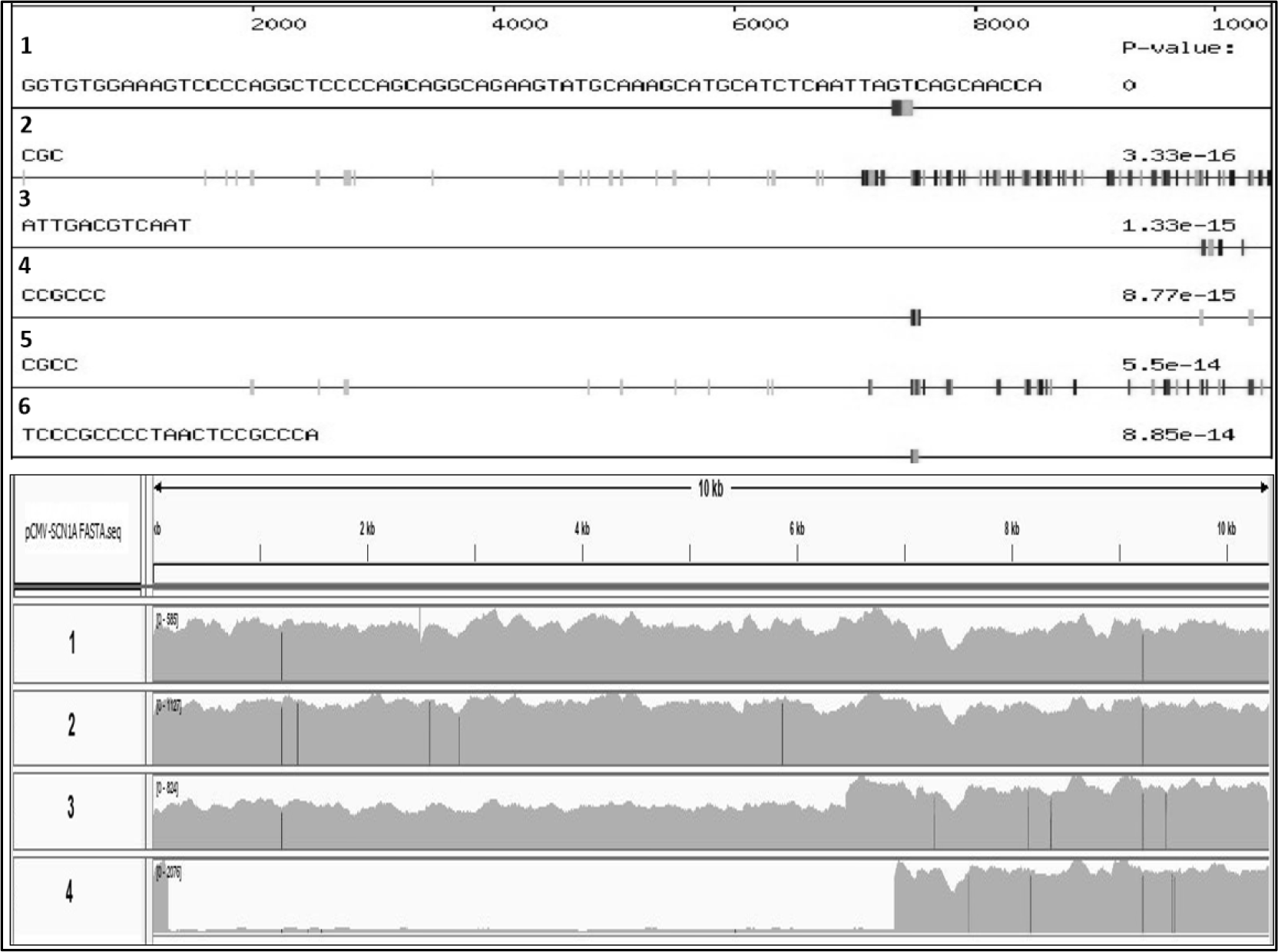
Analysis using the REPFIND program showing the distribution of the repetitive regions in the native plasmid pCMV-*SCN1A*. Black and gray vertical bars show the regions with repetitive units.

### 3.2. Transformation of native plasmids

We observed that the number of colonies formed 72 hours after transformation was slightly higher for NSCC than Stbl2 *E. coli* strains (data not shown). We obtained two types of colonies for both *E. coli* strains: yellow (mature) and white (growing). In both cases, they were stored at 4°C (Figure 2A). The colonies were classified as follows: i) the **subcloning colonies** are yellow, of a large size, grow faster in the first 48 hours, and are isolated from other colonies; ii) the **recombinant colonies** are yellow and grow in clusters, which could lead to plasmid recombination; therefore, they are not candidates for subcloning; iii) **satellite colonies** are those closest to each other, which can be false positives for the presence of the plasmid because they might grow where enzymes secreted by other bacteria have degraded the antibiotic. We rarely observed satellite colonies, but this could explain why some colonies were subcloned and did not grow in liquid LB medium with kanamycin. However, the native plasmid pCMV-*SCN1A* could also be expelled by growing colonies. The optimal time interval of growth for subcloned colonies varied from 48 to 96 hours. We observed that growth efficiency was lower, and the probability of contamination with other microorganisms was higher for more extended incubation periods.

**Figure 2.**
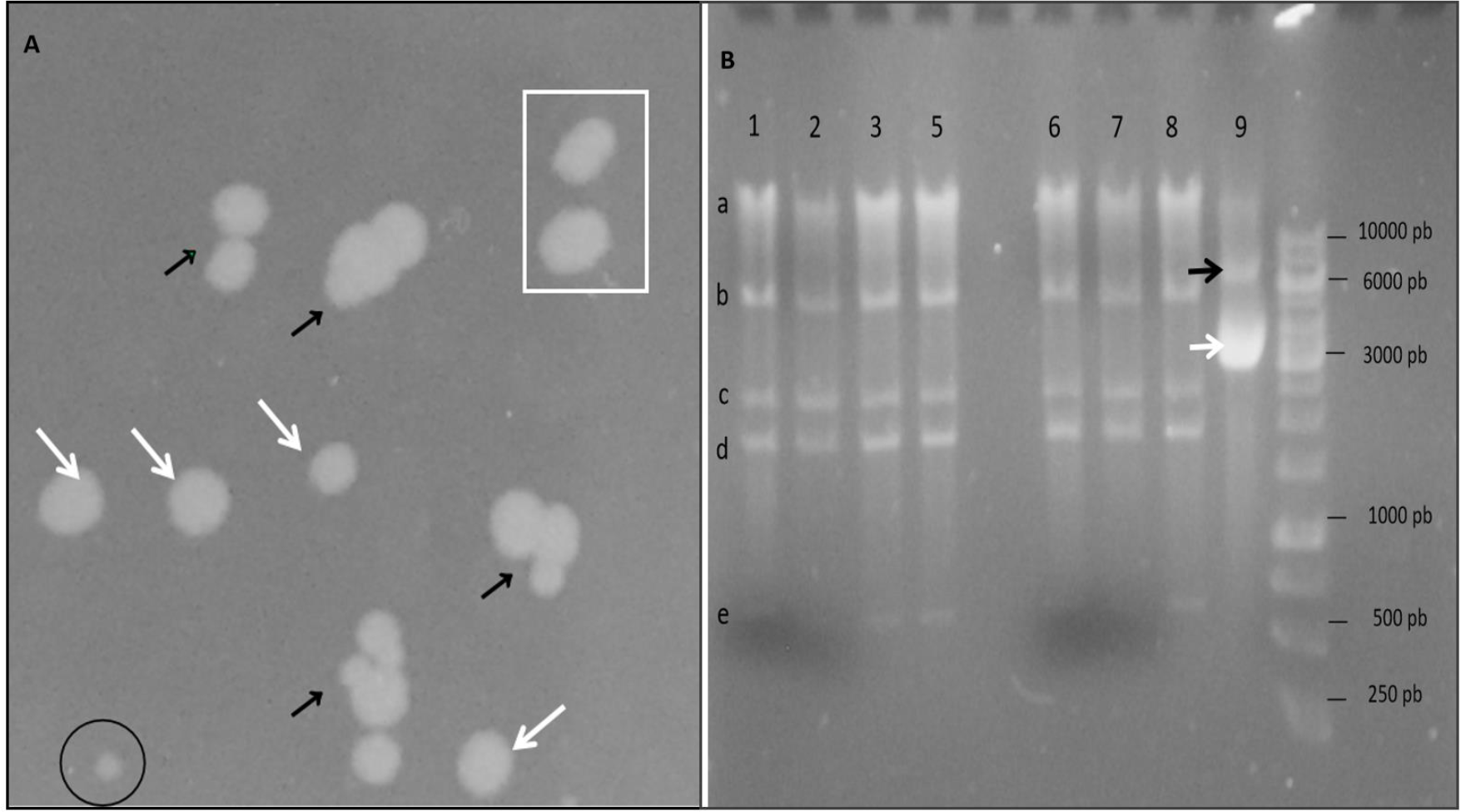
**A)** An image of NEB Stable competent cell (NSCC) colonies transformed with the native plasmid pCMV-*SCN1A* after being grown in solid Luria Bertani (LB) medium with kanamycin for 72 hours. White arrows show colonies that are candidates for subcloning. Black arrows show recombinant colonies. The rectangle shows satellite colonies. The circle contains a small colony. **B)** Enzymatic digestion profile with the enzyme BamHI. Samples 1–9 show plasmids with the expected digestion profile with four fragments: 5.4 kb (b), 2.5 kb (c), 1.9 kb (d), and 620 bp (e). The largest band of 10450 bp (a) corresponds to the native plasmid that the enzyme could not digest. Sample 9 corresponds to a plasmid with structural alterations whose enzymatic digestion shows fragments of 3 and 6 kb (white and black arrows, respectively). The DNA ladder ranges from 250 bp to 10 kb.

We also observed that growing bacterial colonies at 27–30°C slowed down cell metabolism and probably helped maintain the stability of the plasmid in the replication stage. However, we observed the opposite effect at temperatures greater than 30°C (Figure 2A). The subcloning phase of the mini cultures was 48 hours, showing turbidity in the LB medium. Isolated and purified plasmids had a concentration of 40 ng/μl in a final volume of 100 μl.

Analysis of enzymatic digestion assays of mini cultures revealed the expected profile: 5.4 kb, 2.5 kb, 1.9 kb, and 620 bp DNA fragments (Figure 2B). However, recombinant, and non-recombinant plasmids in the same sample were identified in some cell cultures, characterizing mosaicism of the plasmid. This mosaicism was more frequent in cultures from Stb2 cells, probably generated by the instability of the plasmid due to the repetitive regions and resulting in deletions. We also observed that the instability of the plasmid was directly proportional to the volume of the cell culture: at volumes less than 50 ml, this probability decreases. However, there is always greater instability with the Stbl2 cells. One possible explanation is increased cell metabolism at larger volumes of LB culture, and stress leads to plasmid instability.

Subcloning of positive colonies of both bacterial strains was carried out in fractionated volumes of four tubes of 50 ml of LB medium for each colony. After purification of bacterial cultures of both strains, an average concentration of 30 ng/μl plasmid DNA was obtained in a final volume of 1 ml. In some bacterial cultures of both strains, we observed that despite the existence of cell mass, the plasmid DNA concentration was minimal (< 1 ng/μl) or null, this finding being more frequent in the cultures from the Stbl2 cells. This probably occurs because, during cell replication, some bacteria expel the plasmids (large and unstable plasmids) to improve their cellular metabolism and decrease energy expenditure. When the plasmids are expelled in the culture medium, they are degraded (Silva et al., 2012).

Generally, cell cultures are carried out at 37°C; however, at that temperature, even though colonies appeared after 24 hours of incubation, we observed that after enzymatic digestion and sequencing, many plasmids from both strains presented deletions. At 27–30°C, colony growth was observed for both strains after 72 hours. Although the number of colonies was smaller than at 37°C, after enzymatic digestion, fewer deletions were found in the plasmids from the NSCC. This is probably because lower temperatures slow cell growth, causing replication with fewer errors in the plasmid. In addition, there was a higher yield of plasmid DNA in the subcloning stage for both strains when it was purified as soon as possible. This phenomenon occurs because with time, there is an increased probability of the plasmid being expelled.

### 3.3. Site-directed mutagenesis

The annealing temperature for each pair of primers was between 55 and 57°C; however, better results were achieved at 57°C. We used high-fidelity polymerase, which does not generate mutations and does not add any additional nucleotides to the PCR product. The temperature and elongation time depend on the characteristics of the polymerase used. In this case, it was 1 kb per 30 seconds at a temperature of 68°C, so elongation of the 10451 bp parental plasmid would take 5 minutes and 30 seconds. Longer times can generate non-specificity and decrease the plasmid concentration. The PCR products are mutant linear plasmids (MLP) that differ from the parental plasmid because they are linear, not methylated, carry the mutation of interest, and can be converted into a circular shape by recombination. By contrast, a parental plasmid is circular, generated from cell cloning, and has the nucleotide sequence of interest without any mutation and with different degrees of coils (Figure 3A).

**Figure 3.**
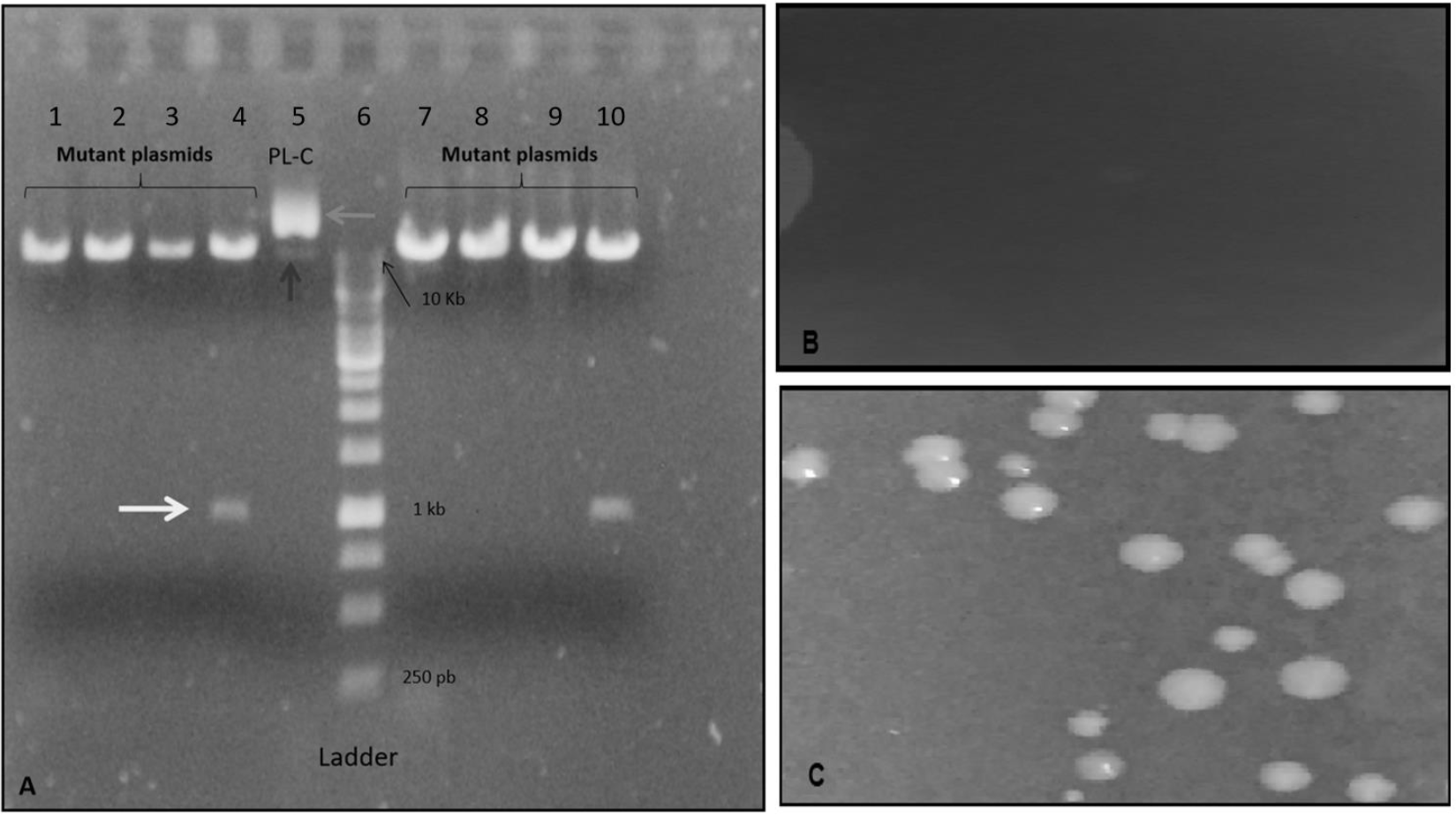
A) Mutant linear plasmids (MLP; 1–4 and 7–10), each of which carries a mutation inserted by site-directed mutagenesis. The parental plasmid (5: PL-C) has two to three conformations because it is circular and adopts different degrees of curl (black and gray arrows). Samples 4 and 10 are MLP that generated a non-specific product (white arrow). The DNA ladder ranges from 250 bp to 10 kb. B and C) Images of NEB Stable competent cells (NSCC) subjected to transformation with methylated and unmethylated parental plasmids, cultured in solid Luria Bertani (LB) medium with kanamycin for 72 hours.

Methylation of the parental plasmid was demonstrated after transformation with both bacterial strains. In both cases, colony growth was not generated for these strains compared with controls (parental plasmid without methylation). The lack of growth of the bacterial strains could be because the promoter regions of the parental plasmid pCMV-SCN1A are silenced by DNA methylation.

The main promoter regions in the plasmid are CMV for the *SCN1A* gene and the promoter region for the *kanR* gene. The former contains the GATC and CCTGG sequences, which are susceptible to methylation; when methylated, they cause the silencing of the *SCN1A* gene that is, the absence of the native Nav1.1 protein. Meanwhile, the methylation of the promoter region of the *kanR* gene results in the absence of resistance to kanamycin (carrier of the parental plasmid), inhibiting cell growth (Figure 3B and C). Therefore, it is crucial to prevent the expression of the parental plasmid, ensuring only the expression of the mutant Nav1.1 protein because the simultaneous expression of the native and mutant Nav1.1 protein would generate erroneous electrophysiological recordings.

When testing the protocol presented here, we generated four types of MLP: pCMV-*SCN1A*_5434, pCMV-*SCN1A*_5329, pCMV-*SCN1A*_5864, and pCMV-*SCN1A*_5177. Mutation 5434 is a missense variant (T>C), which generates a change from tryptophan to arginine at amino acid 1812 of the polypeptide. According to the *in-silico* prediction analysis, this variant is predicted as deleterious by all algorithms; thus, it would likely lead to severe functional changes in the protein. Mutation 5329 is a deletion (delG) that results in a frameshift change, generating a premature stop codon potentially leading to a truncated protein with loss of the C-terminal portion, which is essential for anchoring the protein to the membrane and for regulation. Mutation 5864 is a missense variant (T>C), which generates a change from isoleucine to threonine at amino acid 1955 of the polypeptide. According to the prediction algorithms, the amino acid change is considered benign and predicted not to affect the protein function. Finally, mutation 5177 is a nonsense variant (G>A) that generates a premature stop codon in the pore-forming region, leading to a truncated protein and loss of the C-terminal segment.

Although MLP differs by a single nucleotide, the electrophoretic pattern showed that all four plasmids were the same size. The electrophoretic migration pattern of MLP and the parental plasmid showed differences in migration. This may be because the circular parental plasmid presents different degrees of coiling (Figure 3A).

Some MLP were subjected to *in vitro* recombination reactions that made it possible to generate circular mutant plasmids. These mutant plasmids were used to transform NSCC and Stbl2 cells with the standardized temperature, volume, and purification time parameters described in this paper. As a result, more colonies were observed in the NSCC than in the Stbl2 cells. Furthermore, when transforming the NSCC and Stbl2 cells with MLP without recombination, we observed that the NSCC showed greater growth and more colonies compared with the Stbl2 cells.

This observation led us to hypothesize that the NSCC have a better metabolic ability to induce the incoming MLP to acquire the circular format, thus making it easier for subsequent cloning and leading to an increased number and better growth of the NSCC colonies.

### 3.4. Sequencing plasmids

The sequencing of native plasmids after subcloning with the NSCC revealed that three plasmids had the correct sequence, and 21 plasmids presented mutations (Figure 4). Meanwhile, in the subcloning with the Stbl2 cells, we needed to sequence over 70 plasmids to obtain at least one with the correct sequence. Thus, despite the limitations in plasmid replication in NSCC, our data show that they are more efficient than Stbl2 cells to subclone this type of native plasmid.

**Figure 4.**
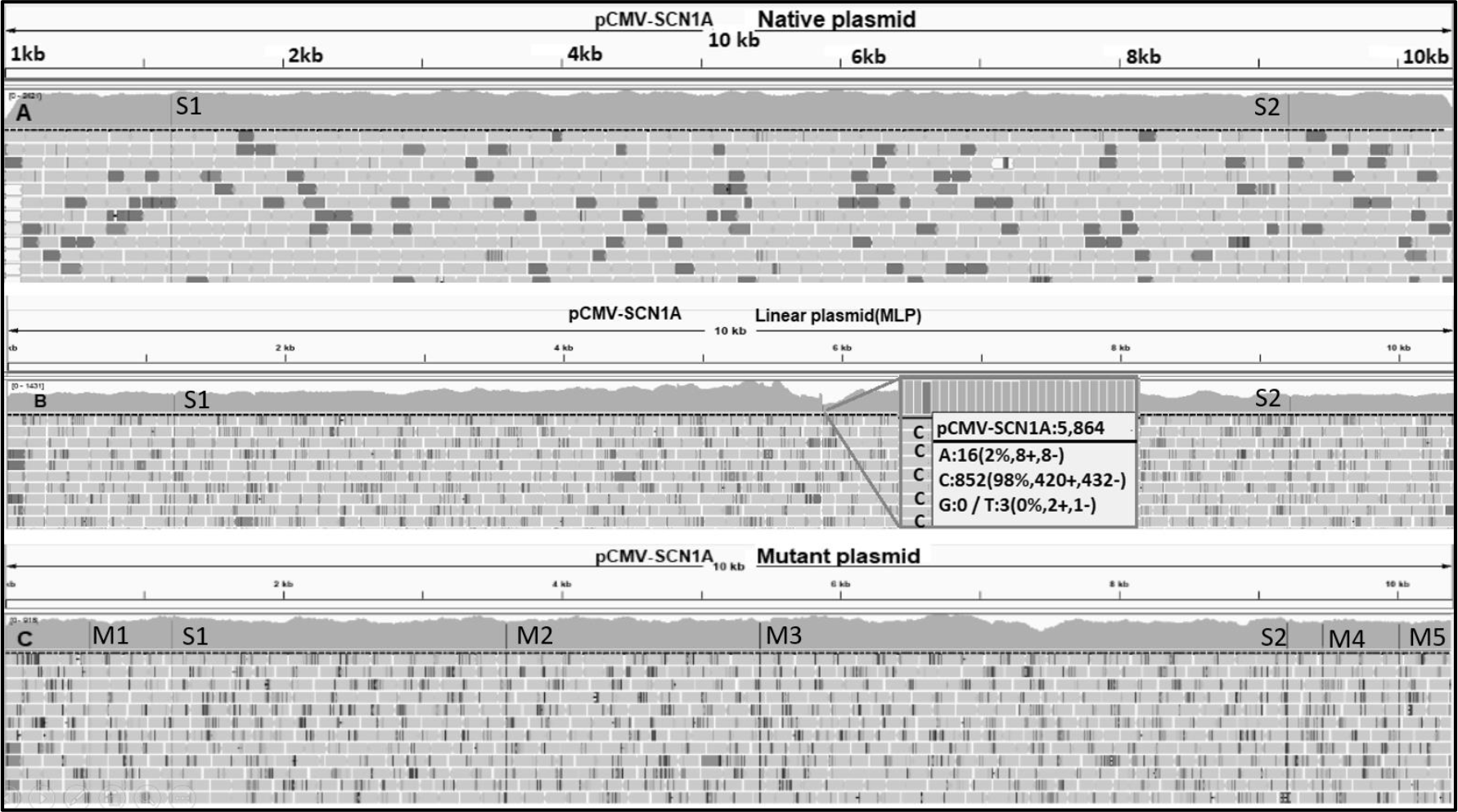
Sequence analysis of the cloned native plasmids. A) A plasmid with two silent mutations (S1 and S2) and a plasmid with the correct sequence. B) Sequencing of site-directed mutagenesis linear plasmids. The square indicates that the 5864 mutation is thymine (3% of reads) for cytosine (98% of reads) substitution. C) A plasmid with point mutations (M): M1-3 are located in the *SCN1A* gene (1-6230 bp). M4-5 located in the vector region (6,231-1045 bp). All these mutations were generated in the subcloning stage. The silent mutations do not affect the expression of the Nav1.1 protein subunit. In other cases, the mutations may have a functional effect if they alter the subunit or some region of interest of the vector.

Regarding MLP, 24 plasmids had the expected sequences, and no unexpected variations were detected. This is because MLP were obtained by PCR and were not cloned in any cell line; thus, the plasmids were not exposed to the metabolic load of the cell during the cloning stage, decreasing the probability of mutation. It is important to indicate that sequencing an MLP before subcloning it into a cell serves as a checkpoint, indicating whether our site-directed mutagenesis experiment resulted in the expected mutation. Indeed, if we only sequenced after plasmid cloning, we could have the following possibilities: 1) find mutations that affect the expression of the gene and the mutation of interest; 2) find only mutations that affect the expression of the gene and the absence of the mutation of interest; 3), find the mutation of interest and mutations that do not affect the expression of the gene of interest because they are located in other plasmid regions that do not involve the gene; and 4) find only the mutation of interest. In cases 1 and 2, the plasmids could not be used for an expression study. In addition, we could not know whether these mutations had come from the mutagenesis reaction or had originated in the cell cloning stage. In cases 3 and 4, the plasmids could be used for expression studies and characterization by cell electrophysiology. Thus, sequencing before cloning allowed us to readily optimize our experiments, saving time and reagents.

After cloning the MLP plasmids (without in vitro recombination) in the Stbl2 cells, little colony growth was achieved, and the plasmids obtained presented deletions. This is probably due to the low efficiency of Stbl2 cells in recirculating MLP, which results in the loss of plasmid fragments or fragments that could be integrated into the bacterial genome. Meanwhile, the mutant plasmids that came from subcloning with NSCC revealed efficient cell growth and only single nucleotide mutations; therefore, the probability of obtaining a desired mutant plasmid was higher. Thus, although the number of colonies was not determined, our results indicate that the metabolic capacity of the NSCC strain is superior because it can better recirculate the plasmid and it is less likely to generate plasmids with unwanted deletions.

The results of sequencing MLP with *in vitro* recombination and cloning in NSCC revealed that the mutant plasmids had single-nucleotide mutations. Indeed, we observed that there is a probability of 1/960 to obtain the plasmid with the desired sequence (one of every 960 transformed NSCC colonies would contain the desired plasmid). Considering Stbl2 cells, this probability is much lower: 1/2880 (one of every 2880 transformed Stbl2 colonies would contain the desired plasmid). Moreover, some of the plasmids obtained from subcloning with Stbl2 cells had deletions. The deletions generated, in most cases, affected a large region of the *SCN1A* gene. We have identified that bacterial strains tend to reduce the size of plasmids in order to save metabolic energy, which is why many deletions of the plasmid are observed in most colonies.

The digestion profiles of BamHI on the plasmid pCMV-*SCN1A* make it possible to determine which cell colonies are potential candidates to be sequenced. The probability of obtaining a colony with a positive BamHI digestion profile is 1/40 for NSCC colonies and 1/120 for Stbl2 colonies (Figure 5). Therefore, in the case of NSCC, of 24 colonies with a positive BamHI profile, only one presented the plasmid with the expected sequence. In the case of Stbl2 colonies, we found that of 80 colonies with a positive BamHI profile, only one of these colonies presented the plasmid with the expected sequence.

**Figure 5.**
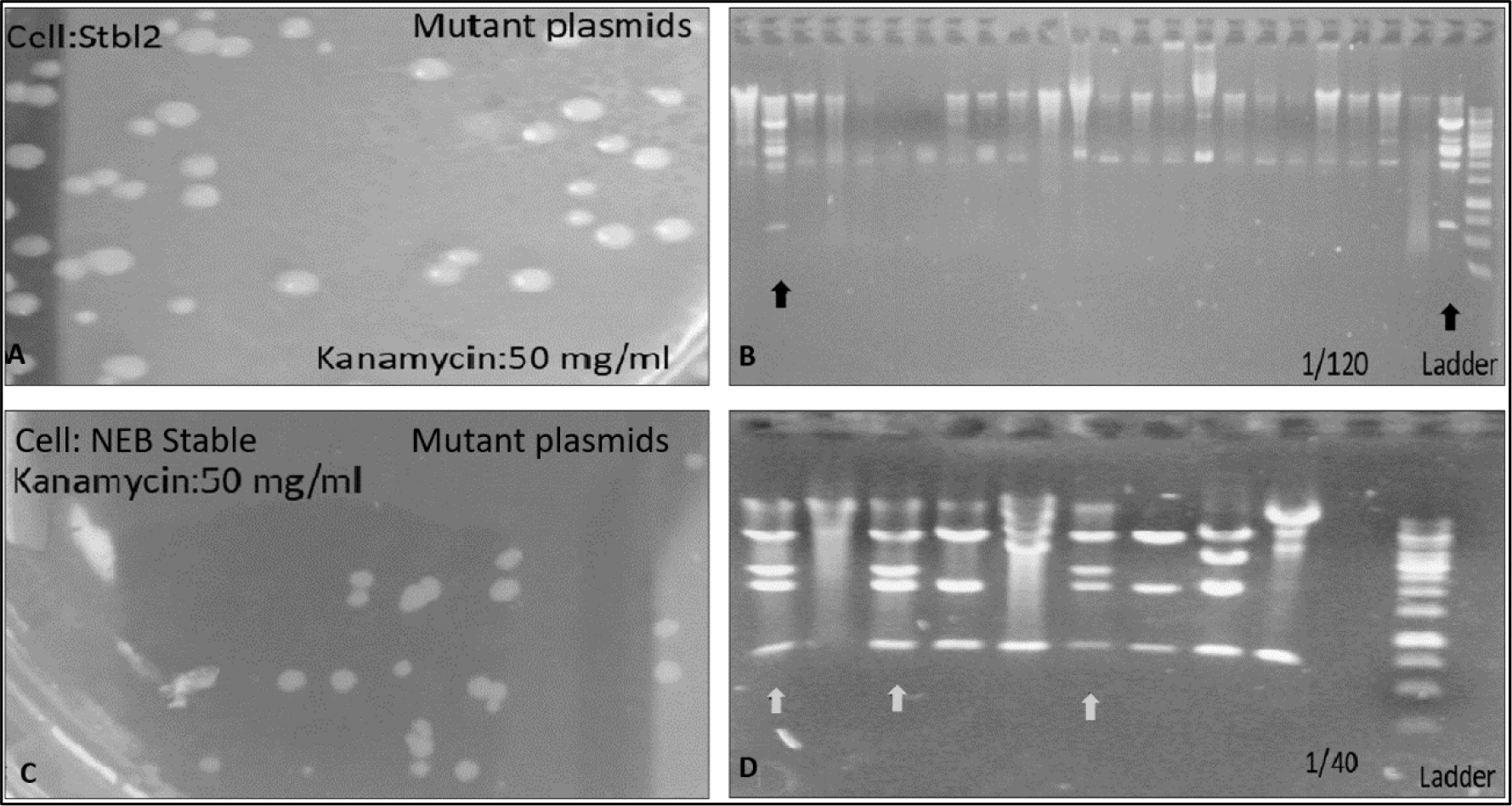
A and C) Post-transformation Stbl2 and NEB Stable cell (NSCC) colonies carrying the same mutant plasmid and growing in solid Luria Bertani (LB) medium. B) Plasmids isolated from Stbl2 cells show different digestion profiles, and only two show the four fragments of interest with an additional 6 kb fragment (black arrows). D) Several of the plasmids isolated from NSCC show the expected digestion profile (gray arrows). In addition, other digestion profiles were identified. The DNA ladder ranges from 250 bp to 10 kb. The samples were run on a 0.8% agarose gel.

## 4. CONCLUSIONS

When cloning highly unstable plasmids encoding sodium channel subunits, point mutations and deletions are frequently generated in the cell subcloning stage. This phenomenon could be due to the high metabolic load of the cells leading to unstable plasmids recombining, generating unexpected variation. Therefore, to clone unstable plasmids, the following factors are critical: 1) use a bacterial strain that can support a high metabolic load; 2) perform cultures in small volumes of LB medium; 3) maintain the temperature between 27 and 30°C, thus promoting a slow growth of bacterial cultures; 4) perform the plasmid purification within 24 hours after culture; and 5) in the case of linear plasmids obtained by site-directed mutagenesis, *in vitro* recombination should be performed before the transformation. Our results have demonstrated that the plasmid pCMV-*SCN1A* presents regions of high instability, leading to the generation of unexpected deletions or point mutations. Furthermore, the use of NSCC combined with the protocol described herein significantly improved the replication of the native pCMV-*SCN1A* and the mutant plasmids created for subsequent functional studies.

## Footnotes

* http://fathmm.biocompute.org.uk

† http://bg.upf.edu/condel/home

‡ http://www.mutationtaster.org

§ http://www.pantherdb.org/tools/csnpScoreForm.jsp

** http://snps.biofold.org/snps-and-go/snps-and-go.html

†† http://mutpred.mutdb.org

‡‡ http://provean.jcvi.org/protein_batch_submit.php?species=human

§§ http://cadd.gs.washington.edu

*** http://genetics.bwh.harvard.edu/pph2/bgi.shtml

††† http://mutationassessor.org/r3

‡‡‡ http://siftdna.org/www/SIFT_pid_subst_all_submit.html

§§§ http://agvgd.hci.utah.edu/agvgd_input.php

**** http://snps.biofold.org/phd-snp/phd-snp.html

## REFERENCES

Al-Allaf, F.A., Tolmachov, O.E., Zambetti, L.P., Tchetchelnitski, V., Mehmet, H., 2013. Remarkable stability of an instability-prone lentiviral vector plasmid in Escherichia coli Stbl3. 3 Biotech 3, 61–70. 10.1007/s13205-012-0070-8

Boucher, H.W., Talbot, G.H., Benjamin, D.K., Bradley, J., Guidos, R.J., Jones, R.N., Murray, B.E., Bonomo, R.A., Gilbert, D., Infectious Diseases Society of America, 2013. 10 x ‘20 Progress--development of new drugs active against gram-negative bacilli: an update from the Infectious Diseases Society of America. Clin. Infect. Dis. Off. Publ. Infect. Dis. Soc. Am. 56, 1685–1694. 10.1093/cid/cit152

Corchero, J.L., Villaverde, A., 1998. Plasmid maintenance in Escherichia coli recombinant cultures is dramatically, steadily, and specifically influenced by features of the encoded proteins. Biotechnol. Bioeng. 58, 625–632. 10.1002/(SICI)1097-0290(19980620)58:6<625::AID-BIT8>3.0.CO;2-K

DeKeyser, J.-M., Thompson, C.H., George, A.L., 2021. Cryptic prokaryotic promoters explain instability of recombinant neuronal sodium channels in bacteria. J. Biol. Chem. 296, 100298. 10.1016/j.jbc.2021.100298

Etchuuya, R., Ito, M., Kitano, S., Shigi, F., Sobue, R., Maeda, S., 2011. Cell-to-cell transformation in Escherichia coli: a novel type of natural transformation involving cell-derived DNA and a putative promoting pheromone. PloS One 6, e16355. 10.1371/journal.pone.0016355

Gonsales, M.C., Montenegro, M.A., Preto, P., Guerreiro, M.M., Coan, A.C., Quast, M.P., Carvalho, B.S., Lopes-Cendes, I., 2019. Multimodal Analysis of SCN1A Missense Variants Improves Interpretation of Clinically Relevant Variants in Dravet Syndrome. Front. Neurol. 0. 10.3389/fneur.2019.00289

Lossin, C., Wang, D.W., Rhodes, T.H., Vanoye, C.G., George, A.L., 2002. Molecular basis of an inherited epilepsy. Neuron 34, 877–884.

Meng, H., Xu, H.-Q., Yu, L., Lin, G.-W., He, N., Su, T., Shi, Y.-W., Li, B., Wang, J., Liu, X.-R., Tang, B., Long, Y.-S., Yi, Y.-H., Liao, W.-P., 2015. The SCN1A Mutation Database: Updating Information and Analysis of the Relationships among Genotype, Functional Alteration, and Phenotype. Hum. Mutat. 36, 573–580. 10.1002/humu.22782

Million-Weaver, S., Camps, M., 2014. Mechanisms of plasmid segregation: Have multicopy plasmids been overlooked? Plasmid 75, 27–36. 10.1016/j.plasmid.2014.07.002

Ridenhour, B.J., Top, E.M., 2016. Plasmid Driven Evolution of Bacteria, in: Kliman, R.M. (Ed.), Encyclopedia of Evolutionary Biology.Academic Press, Oxford, pp. 301–306. 10.1016/B978-0-12-800049-6.00237-7

Sendfeld, F., Selga, E., Scornik, F.S., Pérez, G.J., Mills, N.L., Brugada, R., 2019. Experimental Models of Brugada syndrome. Int. J. Mol. Sci. 20. 10.3390/ijms20092123

Silva, F., Queiroz, J.A., Domingues, F.C., 2012. Evaluating metabolic stress and plasmid stability in plasmid DNA production by Escherichia coli. Biotechnol. Adv. 30, 691–708. 10.1016/j.biotechadv.2011.12.005

Veening, J.-W., Blokesch, M., 2017. Interbacterial predation as a strategy for DNA acquisition in naturally competent bacteria. Nat. Rev. Microbiol. 15, 621–629. 10.1038/nrmicro.2017.66

Wallace, R.H., Scheffer, I.E., Barnett, S., Richards, M., Dibbens, L., Desai, R.R., Lerman-Sagie, T., Lev, D., Mazarib, A., Brand, N., Ben-Zeev, B., Goikhman, I., Singh, R., Kremmidiotis, G., Gardner, A., Sutherland, G.R., George, A.L., Mulley, J.C., Berkovic, S.F., 2001. Neuronal Sodium-Channel α1-Subunit Mutations in Generalized Epilepsy with Febrile Seizures Plus. Am. J. Hum. Genet. 68, 859–865. 10.1086/319516

Yoshida, N., Sato, M., 2009. Plasmid uptake by bacteria: a comparison of methods and efficiencies. Appl. Microbiol. Biotechnol. 83, 791–798. 10.1007/s00253-009-2042-4

